# Hemin Triggers Cytoprotective Lipid Droplet Formation in Pulmonary Microvascular Endothelial Cells

**DOI:** 10.64898/2026.10.29.762875

**Authors:** Alan de Brito Carneiro, Robert W. Wysocki, Xu Zhang, Victor R. Gordeuk, Roberto F. Machado

## Abstract

Sickle cell disease (SCD) is characterized by chronic and episodic hemolysis, inducing endothelial cell dysfunction and death. Lipid droplets (LDs) accumulate intracellularly in response to stress but the role of LDs in hemin-stressed pulmonary microvascular cells remain unexplored. We hypothesized that LD formation is an adaptive response that improves endothelial cell survival during hemin-induced stress. We utilized cultures of human lung microvascular endothelial cells (hLMVECs) to test this hypothesis. Hemin alone was able to induce LD biogenesis in culture. In response to hemin treatment, we found an increase in cellular death by lactate dehydrogenase (LDH) assay when LD biogenesis was inhibited with a small molecule inhibitor of diacylglycerol acyltransferase 1 (DGAT1), A922500. Induction of LDs by treatment with oleic acid (OA) coupled with hemin resulted in improved endothelial cell survival. Hemin treatment was able to activate p-JNK and p-P38 signaling, while treatment with OA alone could not induce either signaling pathway. Hemin treatment combined with DGAT1 inhibition resulted in an increase in endothelial cell secreted IL-6 and IL-8 levels as measured by enzyme linked immunosorbent assay (ELISA), demonstrating an increase in endothelial cell dysfunction and stress. Induction of LD formation with oleic acid (OA) treatment resulted a reduction in secreted IL-6 and IL-8. Mitochondrial reactive oxygen species (ROS) production was decreased by OA treatment, as evidenced by MitoSOXRed assay. These data suggest that hLMVECs are protected from hemin-induced inflammation and death by the presence of LDs. Further studies are needed to explore the Hemin-LD-Cell Death axis for translational therapeutic use.

## Introduction

Sickle cell disease (SCD) is one of the most common monogenic disorders worldwide, with an estimated 515,000 affected newborns in 2021 and approximately 7.7 million individuals living globally with the disease (1). SCD is a group of inherited hemoglobinopathies caused by homozygosity or compound heterozygosity for a single point mutation in the β-globin gene (HBB), resulting in a production of abnormal hemoglobin S (HbS) (2). SCD is associated with multiple acute and chronic complications, including vaso-occlusive crisis (VOC) and acute chest syndrome (ACS), which contribute substantially to disease-related morbidity and mortality (3-6). The pathophysiology of both VOC and ACS is multifactorial, involving complex interactions among sickled erythrocytes, leukocytes, platelets, and the vascular endothelium, as well as inflammatory and thrombotic processes (7). Intravascular hemolysis further contributes to endothelial dysfunction, oxidative stress, and chronic inflammation (3, 8, 9). Hemolysis releases free hemoglobin and heme into the circulation which act as potent damage-associated molecular patterns (DAMPs) that lead to the activation of endothelial cells, oxidative stress responses, and pro-inflammatory signaling pathways in VOC and ACS (9-12). A better understanding of how hemolysis-associated endothelial cell stress is initiated and regulated may help identify new pathways and potential therapeutic targets to mitigate VOC and ACS in patients with SCD.

Free heme activates Toll-like Receptor 4 (TLR4) signaling in endothelial cells; leading to upregulation of adhesion molecules such as P-selectin, E-selectin, Vascular Cell Adhesion Molecule 1 (VCAM-1), and Weibel-Palade body degranulation (13, 14). This pro-inflammatory and pro-adhesive endothelial phenotype increases attachment of sickled erythrocytes, leukocytes, and platelets to the vessel wall thereby facilitating microvascular obstruction and directly contributing to development of VOC (15). Moreover, cell-free hemoglobin and heme contribute to vascular oxidative stress by reducing nitric oxide (NO) bioavailability and stimulating reactive oxygen species (ROS) generation and lipid peroxidation. Heme-induced mitochondrial dysfunction may further exacerbate endothelial oxidative stress by promoting mitochondrial ROS production (3, 13). In the pulmonary microvasculature, endothelial dysfunction and increased vascular permeability, together with inflammatory responses, contribute to the pathophysiology of ACS (16-19).

Lipid droplets (LD) are dynamic intracellular organelles specializing in the storage and regulation of neutral lipids (20). These organelles consist of a hydrophobic core of neutral lipid, primarily triacylglycerols (TAGs) and cholesteryl esters (CEs), surrounded by a phospholipid monolayer associated with proteins involved in lipid metabolism and LD dynamics, including the PAT protein family(21). The PAT family, named after its founding members Perilipin, Adipose differentiation-related protein (ADRP), and Tail-interacting protein of 47 kDa (TIP47) proteins, serve as critical regulators of LD dynamics, acting in lipid storage, lipolysis, and organelle crosstalk (21). The synthesis of LD occurs between two leaflets of the endoplasmic reticulum (ER) membrane where TAGs and CEs are synthesized, predominantly by enzymes such as diacylglycerol acyltransferases (DGAT1/2) and acyl-CoA:cholesterol acyltransferase (ACAT), respectively (20, 22, 23). Beyond their traditional role in lipid storage, LDs are increasingly recognized as regulators of cellular stress response, contributing to lipid homeostasis and protection against lipotoxic and oxidative damage in multiple cell types (23). Metabolic, oxidative, and inflammatory stress induce LD biogenesis in endothelial cells (24, 25). LD accumulation has been associated with conditions such as atherosclerosis, diabetes-related vasculopathy, pulmonary hypertension, and obesity (25-28). However, the role of LD in endothelial response to hemolysis-associated stress is largely unexplored.

Hemin treatment of cultured endothelial cells causes oxidative damage and lipid peroxidation, as well as necroptotic cell death (17, 19). Necroptosis is a mechanism of regulated cell death which is characterized by permeabilization of the plasma membrane through the phosphorylation of Mixed Lineage Kinase Domain-Like Pseudokinase (MLKL) by Receptor-interacting Serine-threonine Protein Kinase 3 (RIPK3) rather than through apoptotic pathways involving caspases (29).. The peroxidation of lipids within the endothelial cells leads to mitochondrial dysfunction, characterized by reduced basal oxygen consumption rate (OCR), increasing proton leak, and decreasing ATP-linked respiration (17). While several mechanisms underlying hemin-induced endothelial injury have been described, the cellular mechanisms that may protect endothelial cells against hemin-induced injury remain unclear.

In sickle cell disease models, the role of LD biogenesis in sickle cell disease models remains limited. We sought to test the hypothesis that LD biogenesis is induced by hemin treatment and that LDs have a positive effect on cell survival in this setting. Our data demonstrates the protective effect of LD biogenesis on hemin-treated lung microvascular endothelial cells.

## Material and Methods

### Cell Culture

Human lung microvascular endothelial cells (hLMVECs) obtained from Lonza Bioscience (CC-2257) were grown at 37° Celsius and in a 5% CO_2_ atmosphere in a cell culture incubator using microvascular endothelial growth media containing supplemental growth factors (Lonza Bioscience, CC-4147) and 5% fetal bovine serum (Gibco, 10438-034) according to the manufacturer’s instructions. Cells underwent a maximum of seven passages in experiments. Prior to experimentation, complete growth media was removed and replaced with basal media (without fetal bovine serum or growth factors) 3 hours before hemin (Sigma-Aldrich, H9039) treatment.

### Lipid Droplet Staining

hLMVECs were seeded overnight onto 24-well culture plates with coverslips at a concentration of 1 x 10^5^ cells/well in 500 μL of endothelial growth media. Cells were incubated for 3 hours in the presence or absence of 10, 20, or 40 μM of hemin, and 5 or 50 nM of the DGAT1 inhibitor A922500 (MedChemExpress, HY-10038), or 10 μM of Oleic acid (OA) (Cayman Chemical, 90260). After 3 hours of incubation, cells were washed with phosphate buffered saline (PBS) solution and fixed in 3.7% formaldehyde at room temperature for 20 minutes. Lipid droplets (LDs) were stained with a solution of 0.18% of Oil Red O (ORO) (Sigma-Aldrich, O-0625) diluted in isopropanol 60% for 2 minutes at room temperature, then washed once with isopropanol 30%, and 5 times with distillated water. Nuclei were stained with 4,6-diamino-2-phenylindole (DAPI) (1 µg/mL) for 10 minutes. The coverslips were mounted using Vectashield antifading mounting medium (Vector Laboratories, H-1000-10). The fluorescence signal was captured by fluorescence microscopy (Nikon Eclipse 80i microscope with camera and NIS-Elements 4.0 software) and quantified using ImageJ software (ImageJ 1.52i).

### RNA Extraction and PCR Analysis

Following 1 hour incubation of 2 x 10^5^ hLMVECs in the presence or absence of hemin 40 μM, A922500 5 μM, the DGAT1 inhibitor. Cells were homogenized in TRIzol (Invitrogen, 15596026) and total RNA was isolated following the manufacturer’s instructions. The cDNA was generated using MultiScribe Reverse Transcriptase kit (ThermoFisher Scientific, 43-688-14) and then used for polymerase chain reaction (PCR) in real-time system using specific primers for ADRP (TaqMan; Hs00160173_m1) and constitutive gene GAPDH (TaqMan; Hs00605340_m1) as endogenous control.

### Cytotoxicity Assay

Following incubation of 5 x 10^5^ hLMVECs in the presence or absence of hemin 40 μM, the DGAT1 inhibitor A922500 5 μM, or OA 10 μM for 24 hours. The supernatants were collected and the lactate dehydrogenase activity (LDH) was measured using the CytoTox 96 Non-Radioactive Cytotoxicity Assay as per the manufacturer’s instructions (Promega, G1780).

### Mitochondrial Reactive Oxygen Species Analysis

hLMVECs were seeded overnight into 96-well culture plates with coverslips at a concentration of 1 × 10^4^ cells/well in 200 μL of endothelial growth media. Cells were treated in the presence of 40 μM of hemin and 10 μM of OA. After 3 hours of incubation, MitoSOXRed (Invitrogen, M36008) reagent was added to cells and incubated for 30 minutes. Cells were then washed 3 times with Hank’s balanced salt solution (HBSS), fixed in 3.7% formaldehyde at room temperature for 20 minutes, the nuclei stained with DAPI, and the fluorescence was analyzed by fluorescence microscopy.

### Cytokine Measurements

All cytokines were quantified by ELISA as recommended by the manufacturer (R&D systems). IL-8 (DY208) and IL-6 (DY206) were measured after incubation of 5 x 10^5^ hLMVECs in the presence or absence of hemin 40 μM, A922500 5 μM, or OA 10 μM for 18 hours.

### SDS-PAGE and Western Blotting

Whole cell lysates from 1 x 10^5^ hLMVECs treated with hemin 40 μM, 5 μM A922500, or OA 10 μM were homogenized with RIPA buffer (ThermoFisher Scientific, 89900) plus protease inhibitor cocktail (ThermoFisher Scientific, PI87786); 30-50 μg homogenates were separated by 4–15% precast polyacrylamide gel (Bio-RAD, 5671084) running on a premixed electrophoresis buffer (Bio-RAD, 1610732) at a constant current of 16 mA. The Precision Plus Protein WesternC Blotting Standards Spectra (Bio-RAD, 1610376) was used as a relative molecular mass standard. After the gel run, the samples were transferred to a polyvinylidene fluoride (PVDF) membrane in a buffer of 25 mM Tris– HCl pH 8.3 and 192 mM glycine, at 190 mA for 90 minutes at 4 °C. The membranes were blocked with a Tris buffered saline (TBS) solution containing 0.1% Tween-20 and 5% bovine serum albumin (BSA) for 1 hour and then incubated for 18 hours with primary antibodies (1:1000) at 4 °C. The membranes were then incubated with a secondary antibody (IgG-HRP, Santa Cruz Biotechnology, CA), followed by detection with the Thermo Scientific SuperSignal ELISA Pico Chemiluminescent Substrate (ThermoFisher Scientific, PI37069) and Bio-RAD ChemiDoc MP Imaging System.

Antibodies used were: Anti-ADRP (sc-377429), Perilipin (sc-390169), phospho-ERK (sc-7383), phospho-JNK (sc-6254), or phospho-p38 (sc-7973) from Santa Cruz Biotechnology, Dallas, TX, USA.

### Genetic Association of Expression Quantitative Loci (eQTL) for *DGAT1* with Clinical Outcomes in Patients with Sickle Cell Disease

Institutional Review Board (IRB) of University of Illinois Chicago (UIC) approved collection of the samples and analysis for the UIC Cohort. IRBs of Albert Einstein College of Medicine, Columbia University, Children’s Hospital of Oakland, Children’s Hospital Pittsburgh, Hammersmith Hospital London, Howard University, Johns Hopkins University, NHLBI, University of Colorado, and UIC approved collection of samples and analysis for the Treatment of Pulmonary Hypertension and Sickle Cell Disease with Sildenafil Therapy (Walk-PHaSST) cohort. For further details see Zhang *et al.*, 2023 (30).

Clinical outcomes were obtained from patients in their usual state of health. Array genotyping of the patient cohorts and genotype imputation to 1000 genomes phase 3 data were described previously (30). Bulk single tissue eQTL for *DGAT1* were queried from Genotype-Tissue Expression (GTEx) project version 10. Single nucleotide polymorphism (SNPs) with minor allele frequency >0.1 and imputation quality r^2^ >0.8 across both cohorts were analyzed, resulting 204 tested SNPs. Genetic associations with pulmonary vascular resistance (PVR) and acute chest syndrome (ACS) used linear regression and logistic linear regression, respectively, with an additive genetic mode and controlling for age, gender, Hb genotype severity (SS/Sβ^0^ versus SC/ Sβ^+^), hydroxyurea treatment, and population stratification. Meta analysis used inverse variance weighting (31). Significance was determined by a false discovery rate (FDR) of 0.05.

## Results

### Hemin Induces Lipid Droplet Biogenesis in Human Lung Microvascular Endothelial Cells

We sought to model the effect of intravascular hemolysis on lipid droplet (LD) biogenesis by utilizing the *in vitro* hemin model of hemolytic injury. Human lung microvascular endothelial cells (hLMVECs) were treated with increasing concentrations of hemin, the oxidized form of heme released during intravascular hemolysis, and LD production was measured by oil red o (ORO) staining. After 3 hours of incubation, we noted an increase in ORO staining when hLMVECs were treated with increasing concentrations of hemin, suggesting an increase in LD formation (Figure 1A).

**Figure 1:**
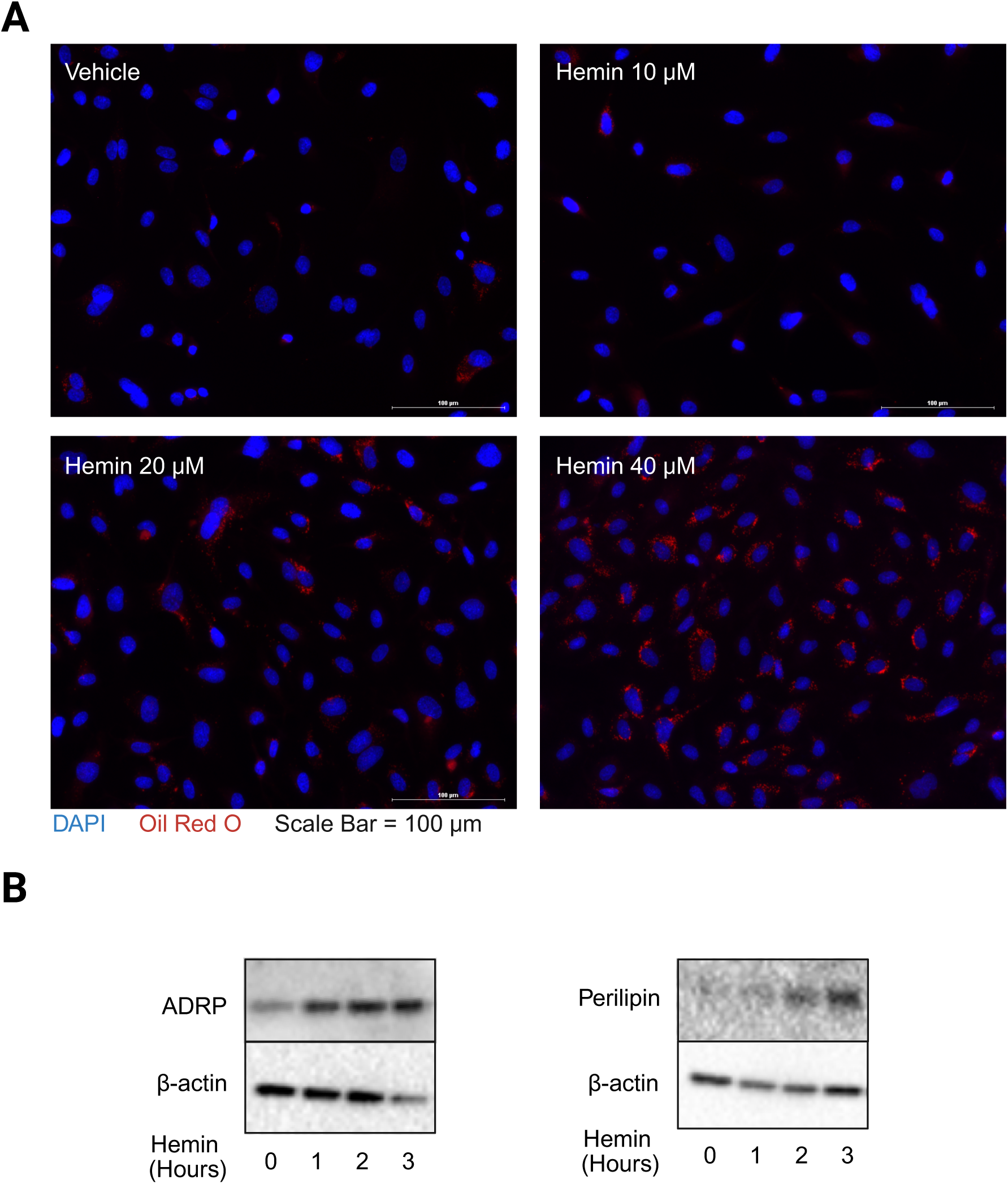
Hemin induces Lipid Droplet formation in human lung microvascular endothelial cells (hLMVECs). **(A)** hLMVECs were incubated in the presence of 10, 20, or 40 μM of hemin or vehicle control. Oil Red O (ORO) staining was used as a marker of lipid droplet formation within the cells. **(B)** hLMVEC were treated with 40 μM hemin along a time-course from 0 to 3 hours. Western blot of whole cell lysates were probed for ADRP and Perilipin protein levels.

LD biogenesis involves multiple structural proteins, two of which are adipose differentiation-related protein (ADRP) also known as perilipin-2 and perilipin (21). To confirm that the ORO staining increase is from LD biogenesis, we treated hLMVEC with hemin over a 1–3-hour time-course. Western blot probed for ADRP and perilipin protein levels confirmed an increase in these LD structural proteins in response to hemin treatment (Figure 1B). These data further suggest that hemin inducing LD formation in hLMVECs.

### Lipid Droplets Mitigate Hemin-induced Cell Death

Inhibition of DGAT1 with a potent small molecule inhibitor, A922500, resulted in a decrease in hemin-induced LD formation both qualitatively and quantitatively in hLMVECs (Figure 2A-B) (32). Quantitative PCR shows that mRNA for ADRP is decreased when DGAT1 is inhibited by A922500 treatment is combined with one hour treatment with hemin (Supplemental Figure 1).

**Figure 2:**
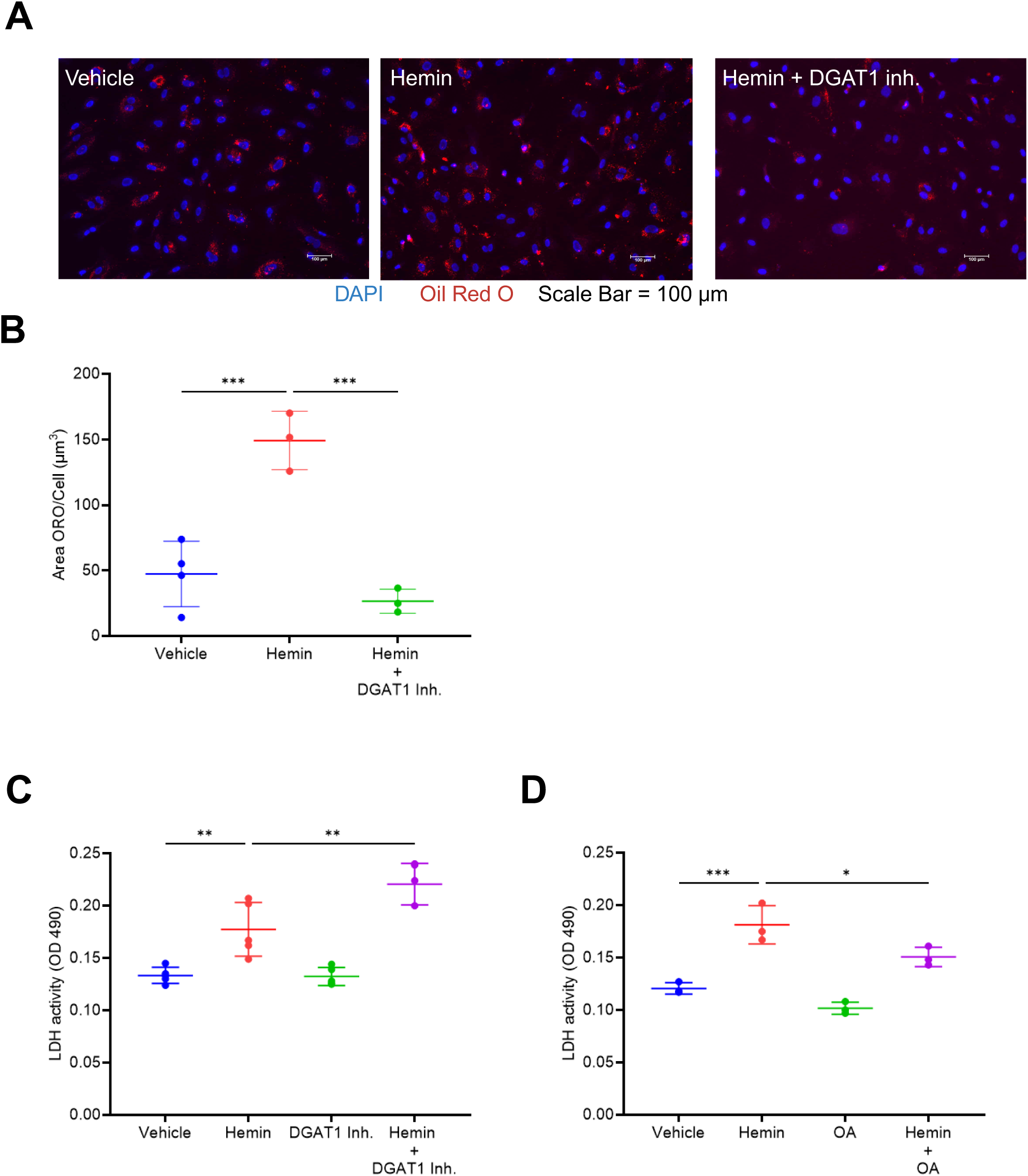
Lipid droplets attenuate hemin-induced cell death. **(A)** hLMVEC were treated for 3 hours with 40 μM hemin, and/or 5 nM A922500, a DGAT1 inhibitor, with representative images of immunofluorescence shown. **(B)** Quantified Oil Red O (ORO) area per cell area (ImageJ 1.52i), n=3-4. **(C-D)** Cell death was measured by a lactate dehydrogenase (LDH) activity colorimetric assay kit for cells treated with hemin, A922500, oleic acid (OA), or combination. (n=3-4).**** indicates p-value ≤ 0.0001; *** indicates p-value ≤ 0.001; ** indicates p-value ≤ 0.01; * indicates p-value ≤ 0.05.

Lactate dehydrogenase (LDH) is released from multiple cell types suffering from membrane damage, serving as a reliable marker of cell death (33). hLMVECs showed a significant increase in LDH activity when DGAT1 inhibition is combined with hemin treatment versus with hemin treatment alone, indicative of LD protection against hemin-induced cell death (Figure 2C).

Oleic acid (OA) has been shown to be an inducer of LD biogenesis (34). We treated hLMVECs with hemin or with hemin and OA concurrently to determine if an increase in LD biogenesis could attenuate hemin-induced cell death. We saw a significant decrease in LDH activity when OA was combined with hemin treatment (Figure 2D).

### Exogenously Stimulated Lipid Droplet Biogenesis Does Not Robustly Activate MAPKs JNK, P38, nor ERK

Mitogen-activated protein (MAP) kinase comprises a family of protein-serine/threonine kinases that control intracellular events like growth, division, differentiation, and survival (35). Hemin has been shown to trigger the MAPK pathway including ERK (36). LD formation also involves MAPK signaling in multiple cell types, including immune cells such as monocytes, which activate p38 and JNK (37). We sought to confirm that hemin triggers the MAPK pathway and whether these pathways might be involved in endothelial cell LD formation. hLMVECs cells were incubated with a concentration range of 5-40 μM of hemin. At the dose of 40 μM of hemin, p38 and JNK demonstrated an increase in phosphorylated protein levels (Figure 3A). When exposed to OA, we noted no change in p-JNK, p-P38, or p-ERK protein levels compared to baseline (Figure 3A).

**Figure 3:**
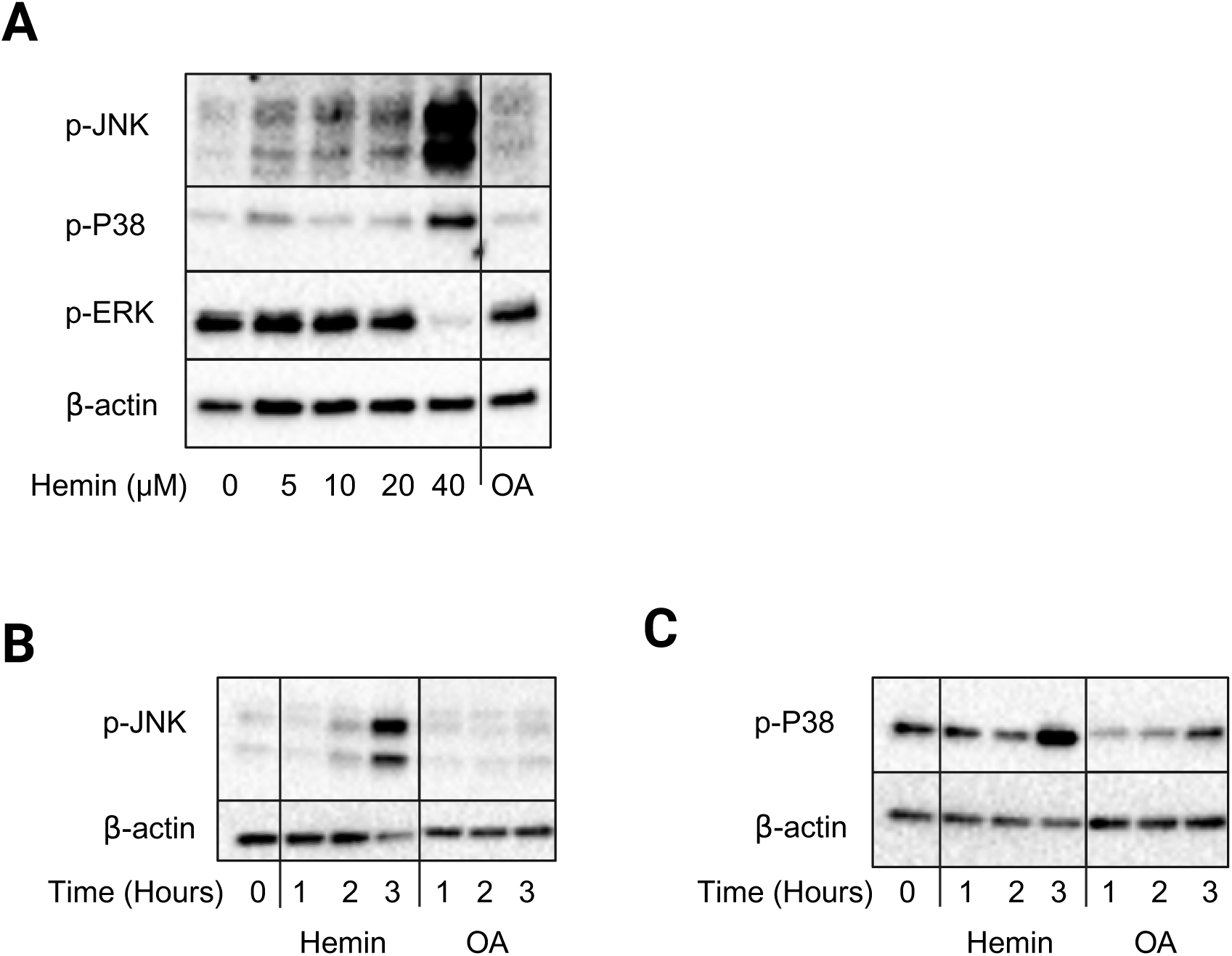
Oleic acid induction of Lipid Droplets is not dependent on activation of the MAPK pathway in human lung microvascular endothelial cells (hLMVECs). **(A)** Western blot of hLMVEC whole cell lysates probing phosphorylated JNK, P38, and ERK over a dose-response of hemin or oleic acid (OA). **(B-C)** hLMVECs were incubated with 40 μM hemin for a 3-hour time-course. Western blot probed for phosphorylation of JNK and phosphorylation of P38.

Because phospho-JNK and phospho-p38 were induced by hemin treatment, we performed a time-course treatment with hemin and OA to determine the timing of the phosphorylation of these enzymes. hLMVECs cells were treated for 1-3-hours with hemin 40 μM or OA. Both JNK and P38 showed increasing phosphorylation over the time course of hemin stimulation. OA stimulation over the 3-hour course did not result in an increase in p-JNK levels but did increase p-P38 across the time course (Figure 3B-3C).

### LD biogenesis Mitigates Inflammatory Cytokine Production, and Mitochondrial ROS Production Induced by Hemin

Because endothelial stress is associated with increased production of IL-6 and IL-8, we next examined whether hemin stimulation induces release of these proinflammatory cytokines in hLMVECs (38, 39). Treatment of hLMVECs with 40 μM hemin resulted in an increase in IL-8 release, further increased by co-treatment with A922500, a DGAT1 inhibitor (Figure 4A). OA co-treatment with hemin was able to significantly decrease IL-8 release when compared with hemin alone, suggesting a diminished cellular stress response (Figure 4B).

**Figure 4:**
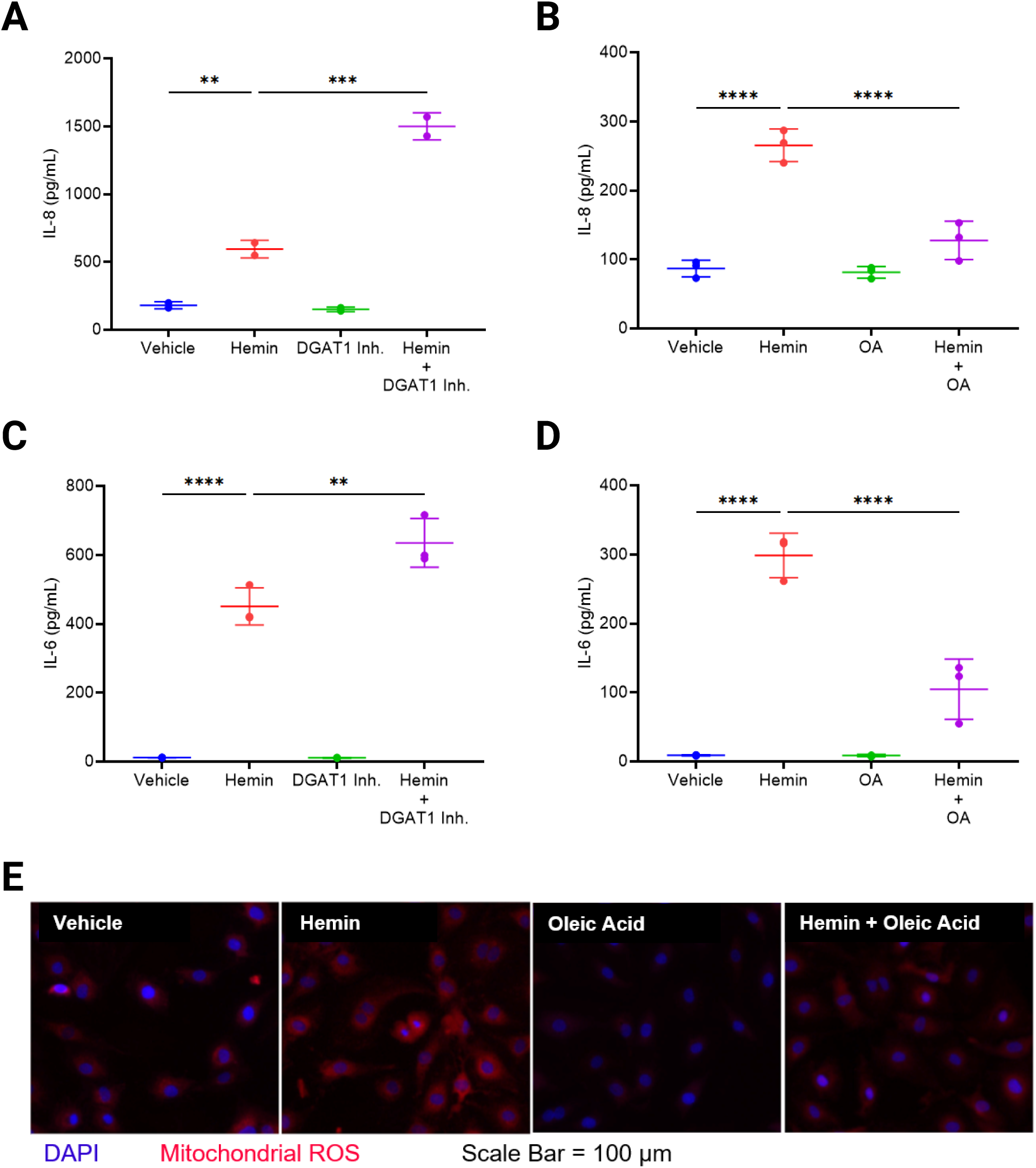
Lipid Droplet biogenesis decreases the pro-inflammatory cytokine production and mitochondrial ROS production that is induced by hemin. Human lung microvascular endothelial cells (hLMVECs) were incubated in the presence or absence of 40 μM hemin, 5 nM of A922500, a DGAT1 inhibitor, or 10 μM Oleic acid (OA). After 24 hours, the supernatants were collected. **(A-B)** IL-8 levels of the supernatant were measured by ELISA, n=2-3. **(C-D)** IL-6 levels of the supernatant were measured by ELISA, n=3. **(E)** Cells were incubated in the presence or absence of 40 μM of hemin or 10 uM of OA for 3 hours. MitoSOX Assay was performed according to manufacturer’s instructions and representative images shown. **** indicates p-value ≤ 0.0001; *** indicates p-value ≤ 0.001; ** indicates p-value ≤ 0.01; * indicates p-value ≤ 0.05.

Similarly, hemin treatment induced IL-6 secretion, which was significantly augmented by inhibition of DGAT1 (Figure 4C). The effect of OA co-treatment with hemin was able to decrease excreted IL-6 levels in hLMVEC supernatant (Figure 4D).

At basal levels, mitochondrial ROS have an important role in endothelial cell homeostasis by regulating vascular tone and angiogenesis (40-42). LD can act as key regulators of cellular redox balance through the induction or inhibition of ROS generation (23). When hLMVECs were incubated with hemin, a strong increase in mitochondrial ROS was seen. This robust increase in mitochondrial ROS production was attenuated by co-treatment with hemin and OA (Figure 4E).

### eQTL for *DGAT1* are Associated with Pulmonary Vascular Resistance and Acute Chest Syndrome Risk in Patient Cohorts

LD biogenesis is partially driven by the enzyme diacylglycerol O-Acyltransferase 1 (DGAT1), which catalyzes the conversion of diacylglycerol and fatty acyl-CoA into triacylglycerol (22, 24). Population studies conducted in our laboratory of the University of Illinois Chicago (UIC) and the Walk-Treatment of Pulmonary Hypertension and Sickle Cell Disease with Sildenafil Therapy (Walk-PHaSST Cohort) revealed two eQTL for *DGAT1*, associated with PVR and ACS clinical outcomes (30, 43, 44). The G allele of SNP rs6986110 was associated with an increase in *DGAT1* expression in multiple tissues, including lung tissue, and with a reduction in PVR in both the Walk-PHaSST cohort (β = -1.47 ± 0.641, *P* = 0.043, *N* = 17) and the UIC cohort (β = -1.54 ± 0.422, *P* = 0.0045, *N* = 16), with a combined β = -1.52 ± 0.353, *P* = 1.6 x 10^-5^ (Supplemental Figure 1A; Supplemental Table 1). The G allele of another SNP rs62530366 was associated with decreased *DGAT1* expression in left ventricular heart tissue and with a reduced risk of ACS in the Walk-PHaSST cohort (β = -0.637 ± 0.169, P = 1.4 × 10⁻⁴, N = 524), but less significant in the UIC cohort (β = -0.266 ± 0.195, P = 0.17, N = 381), with a combined β = -0.477 ± 0.128, P = 1.9 × 10⁻⁴ (Supplemental Figure 1B; Supplemental Table 1).

## Discussion

In this study, we demonstrate that hemin is a robust promoter of LD formation in endothelial cells, and that LDs serve a protective role in these cells. Pharmacological inhibition of DGAT1 suppressed LD synthesis, enhanced inflammatory cytokine release, and increased hemin-induced cell death. Conversely, stimulation of lipid droplet biogenesis exogenously by OA treatment had a protective effect on cells in response to hemin exposure and reduced inflammatory cytokine excretion. These data identify endothelial LD formation as an adaptive response that mitigates hemin-induced injury.

Endothelial cells form LDs when exposed to various forms of extracellular stress including TNF treatment, hypoxic environments, LPS administration, and high fat diet in animal models (45-47). These studies demonstrate that inflammation and endothelial cell stress play a large role in LD formation. Our data supports the conclusion that the increase in ORO staining we see is from LDs rather than nonspecific lipid accumulation. The increased ORO staining in response to hemin treatment is coupled with an increase in protein levels of ADRP and perilipin, two proteins supporting the structure of LDs specifically.

Hemin has been used as a model system for intravascular hemolysis in both *in vitro* and *in vivo* studies, as chronic hemolysis has been shown to increase VOC and ACS events in patients with sickle cell disease (11, 16, 48). Data from our group and others has demonstrated that hemin treatment of endothelial cells induces lipid peroxidation resulting in mitochondrial dysfunction, increases release of IL-6 and IL-8, and causes cell death (17, 49). In the present study, we link the dysfunctional mitochondrial ROS production, cytokine release, and cell death to LD biogenesis induced by hemin. LDs act as suppressors of inflammatory cytokine release and mitigators of cell death. When LD were blocked pharmacologically or increased through lipid treatment induction, we saw a significant effect on these inflammatory and cell death outputs. Similarly, in adipocytes, fibroblasts, and multiple cancer cell lines, genetic knockout or small molecule inhibition of DGAT1 resulted in mitochondrial dysfunction and cellular death (50-52). The robust accumulation of LDs, through an enhancement of ORO staining and increase of LD structural proteins seen in our results, may represent an adaptive response to limit the damage caused by lipid oxidation. Thus, LDs appear to preserve endothelial cell homeostasis in response to free hemin.

The effect of hemin treatment versus oleic acid (OA) treatment on the MAPK signaling pathway suggests that lipid droplet biogenesis is not completely regulated by MAPK. Microglia cells were shown to require p-P38 activation to form lipid droplets in response to LPS treatment, but not to activate JNK (53). Monocytes, however, required both p-P38 and p-JNK for lipid droplet formation in response to arachidonic acid stimulation (37). Hemin, in other models, has been shown to induce p-P38 activation, resulting in endothelial barrier dysfunction (36, 54). Our data suggests that MAPK is not required for LD formation in human pulmonary microvascular cells through an OA stimulus but might be necessary for hemin-induced LD formation, highlighting different pathways for LD biogenesis.

Hemin induction of IL-6 and IL-8 has been described previously through its activation of endothelial cell TLR4 signaling (15, 49). We found that inhibition of LD biogenesis via a small molecule inhibitor of DGAT1 was able to increase inflammatory cytokine production in human pulmonary microvascular cells, while OA treatment was able to partially suppress the inflammatory cascade caused by hemin treatment. Mitochondrial ROS production was also impaired when LDs were formed by OA treatment. Together, these results suggest a protective effect of LDs on endothelial damage signals and ROS production. LDs are known to play a role in intracellular ROS homeostasis (55). TLR4 signaling depends, in part, on mitochondrial ROS production, which is partially suppressed by LDs in our model. TLR4 signaling itself has also been linked to TAG accumulation and DGAT2 activation, both necessary for LD biogenesis (56). Supporting our findings, a recent study has demonstrated that DGAT-1 dependent LD biogenesis protects cardiac endothelial cells from lipotoxic stress, while inhibition of LD synthesis exacerbates mitochondrial dysfunction, lipid peroxidation, and cell death, suggesting LDs as an adaptative cytoprotective mechanism in the cardiovascular system (57). In this study, OA was utilized in higher doses than our study leading to the negative effects of OA on cellular health, suggesting dose-dependency of the protective effects of OA. Genetic manipulation of the DGAT1/2 pathway leading to over-expression and functional knock-down would lend more credibility to the effects demonstrated here, as small molecule inhibitors and OA likely have off-target effects.

Our genetic association results suggest that DGAT1 variants and protein levels influence ACS recurrence risk as well as ACS mortality risk. This result suggests LD play a role in the pathogenesis of ACS. The question, however, is whether endothelial cells or other cell types also require LD biogenesis to combat the effects of chronic and acute intravascular hemolysis. The presence of lower levels of DGAT1 expression in PBMC being protective is interesting, suggesting that LD biogenesis is not always protective in all cells in response to intravascular hemolysis. This would be specifically relevant for T lymphocytes, where LDs were recently shown to be pro-death in an inflammatory model of rheumatoid arthritis, suggesting increasing levels of LD are not beneficial for all cell types as they are in our endothelial cell compartment (58).

The current study demonstrates a role for LDs in the pathogenesis of hemin-induced cellular injury. A limitation of our narrow study is the absence of mechanistic exploration in pulmonary microvascular cells, as has been done in tumor models examining lipid droplets (59). Genetic manipulation of the LD biogenesis pathway in microvascular endothelial cells could identify steps of the pathway amenable to therapeutic targeting. Generalizing our *in vitro* study to *in vivo* effects of lipid droplet induction in sickle cell disease animal models is difficult to speculate. Although we hypothesize that LD would improve endothelial cell survival because of intravascular hemolysis, we have not tested whether LD biogenesis would be able to overcome chronic or acute increases in free hemin products in ACS or VOC models. We do, however, believe that the work presented will form the groundwork for further exploration of LD physiology in models outside of tumorigenesis and atherosclerosis. Our genetic data from our patient cohorts demonstrate an association of PVR and ACS risk with DGAT1 SNPs. The conflicting genetic data, increased DGAT1 expression in lung tissue associated with improved PVR but decreased DGAT1 expression in heart tissue associated with improved ACS risk, suggests that LD may have opposing benefits depending on the tissue in which they are formed. These data further suggest that lipid droplet (LD) pathways influence patient outcomes, although the protective versus detrimental effects of LDs remain incompletely understood. With further research and mechanistic understanding, LDs might be a useful tool in the future to alleviate the risk of ACS or VOC in patients with sickle cell disease.

In conclusion, the *in vitro* data of this study strongly suggests that LD biogenesis represents an adaptative endothelial response in lung microvascular cells to mitigate hemin-induced cytotoxicity. Hemin can induce LD biogenesis, and this LD biogenesis is enough to stave off cell death as evidenced by DGAT1 small molecule inhibition resulting in an increase of hemin-induced cell death. OA can mitigate cell death from hemin and improve inflammatory signals arising from the endothelial cells. Further mechanistic and translational studies are necessary before the full therapeutic possibilities of LDs are understood in SCD and endothelial cells.

