## Supplemental Table 1 for "Hemin Triggers Cytoprotective Lipid Droplet Formation in Pulmonary Microvascular Endothelial Cells"

| SNP | Position (GRCh38) | Reference Allele | Effect Allele | DGAT1 Gene Expression in GTEx v10 | | |
|---|---|---|---|---|---|---|
| | | | | Tissue | Normalized Effect Size (NES) | P-value |
| rs6986110 | 8:75,432,123 | A | G | Lung | 0.12 | $4.0 \times 10^{-8}$ |
| rs6986110 | 8:75,432,123 | A | G | Heart - Left Ventricle | 0.13 | $1.90 \times 10^{-8}$ |
| rs6986110 | 8:75,432,123 | A | G | Whole Blood | 0.13 | $3.70 \times 10^{-11}$ |
| rs6986110 | 8:75,432,123 | A | G | Liver | 0.22 | $2.00 \times 10^{-5}$ |
| rs62530366 | 8:144,312,370 | A | G | Heart - Left Ventricle | -0.12 | $3.4 \times 10^{-5}$ |

| SNP | Clinical Trait | Walk-PHaSST | | | | UIC | | | | Combined | |
|---|---|---|---|---|---|---|---|---|---|---|---|
| | | Effect Allele Frequency | β (SE) | P-value | N | Effect Allele Frequency | β (SE) | P-value | N | β (SE) | P-value |
| rs6986110 | Pulmonary Vascular Resistance (PVR) | 0.62 | -1.47 (0.641) | 0.043 | 17 | 0.68 | -1.54 (0.422) | 0.0045 | 16 | -1.52 (0.353) | $1.6 \times 10^{-5}$ |
| rs62530366 | Acute Chest Syndrome (ACS) Risk | 0.22 | -0.637 (0.169) | 0.00014 | 524 | 0.2 | -0.266 (0.195) | 0.17 | 381 | -0.477 (0.128) | 0.00019 |
