## Supplemental Figures 1 and 2 for "Hemin Triggers Cytoprotective Lipid Droplet Formation in Pulmonary Microvascular Endothelial Cells"

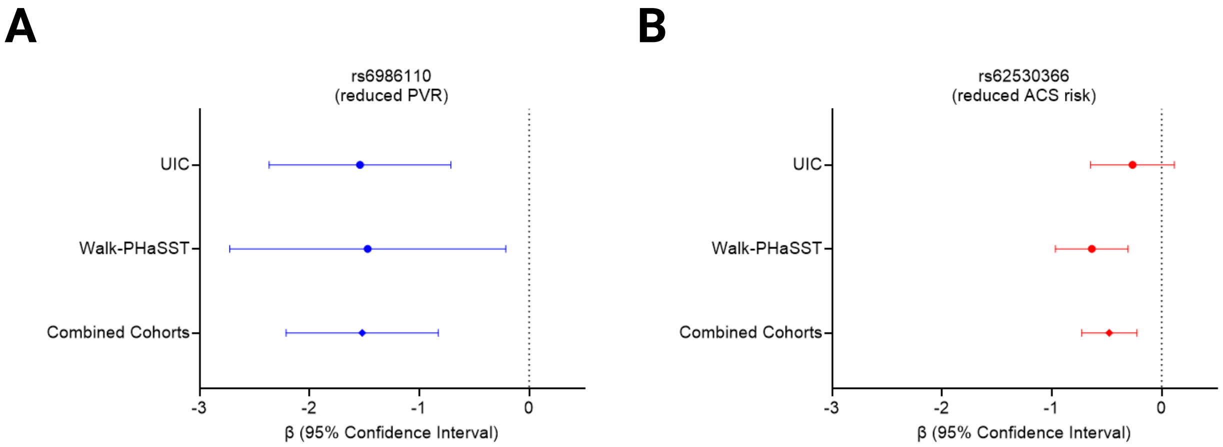


**Supplemental Figure 1: DGAT1 Genetic Variants are Associated with Pulmonary Vascular Resistance and Acute Chest Syndrome Risk in Patient Cohorts. (A)** Forest plot of patient cohort data for DGAT1 variant rs6986110 demonstrating Beta Coefficient (β) with 95% Confidence Interval (CI) for the Walk-PHaSST and UIC Cohorts. **(B)** Forest plot of patient cohort data for DGAT1 variant rs62530366 demonstrating Beta Coefficient (β) with 95% Confidence Interval (CI) for the Walk-PHaSST and UIC Cohorts.

**A**

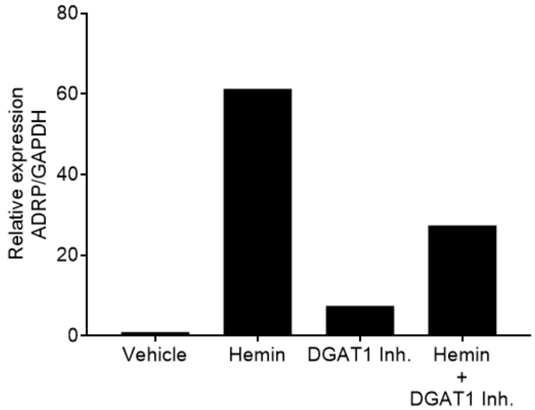


**Supplemental Figure 2: Inhibition of DGAT1 impairs hemin-induced Lipid Droplet biogenesis.** qPCR analysis from whole cell lysates of human lung microvascular endothelial cells (hLMVECs) of ADRP mRNA after 1 hour of incubation 40 μM hemin, or 5 nM A922500, a DGAT1 inhibitor. GAPDH mRNA expression was used as housekeeping control.
